# Mechanistic assessment of eDNA passive samplers: a case study with invasive freshwater bivalves

**DOI:** 10.64898/2026.08.07.743527

**Authors:** Anish Kirtane, Alexandra A.-T. Weber

## Abstract

Passive sampling is the deployment of a collection material in the environment to continuously capture environmental DNA (eDNA) over time, offering the potential to integrate biodiversity signals while reducing the need for repeated active water collection. However, the mechanisms governing eDNA capture and retention on passive samplers remain poorly understood, limiting the interpretation of passive eDNA signals and their broader application. Here, we investigated the mechanistic performance of glass fibre passive samplers using controlled mesocosm experiments with three invasive freshwater bivalves: zebra mussels (*Dreissena polymorpha*), quagga mussels (*Dreissena bugensis*), and Asian clams (*Corbicula fluminea*). Specifically, we quantified eDNA accumulation dynamics, evaluated the contribution of different eDNA states, tested the persistence of captured eDNA, and compared passive sampler signals with conventional active sampling. Passive samplers rapidly accumulated target eDNA within hours of deployment, after which concentrations either plateaued or continued to increase depending on species. Sequential transfer of passive samplers between mesocosms containing different species showed that previously captured eDNA declined while new target eDNA accumulated to concentrations comparable to freshly deployed samplers, demonstrating continual turnover rather than permanent retention. Dissolved eDNA showed little evidence of accumulation beyond the concentration retained in the pore water within the membrane, suggesting that it is unlikely to be the dominant contributor to long-term passive sampler signals. Instead, the observed variability among replicate samplers, together with the physical properties of glass fibre membranes, suggests that membrane-bound and particulate eDNA are the primary contributors to passive eDNA capture. Collectively, these findings support a model in which glass fibre passive sampler signals reflect a dynamic equilibrium between ongoing eDNA capture and concurrent loss processes rather than cumulative accumulation over time. This mechanistic framework provides a foundation for interpreting passive eDNA data and informs the future development of passive sampling materials, deployment strategies, and biodiversity monitoring applications.

## Introduction

Environmental DNA (eDNA) analysis has reshaped how biodiversity is monitored, with applications ranging from broad community surveys to the early detection of invasive species (Deiner et al., 2017; Rishan et al., 2023). At its core, eDNA analysis depends on how effectively genetic material is captured from the environment, most commonly from aquatic systems (Kirtane et al., 2024; Takahashi et al., 2023). The most widely used approach is active sampling, where a discrete water sample is collected and passed through a 0.2–10 µm pore size filter membrane, enabling subsequent analysis of the species composition and/or concentration of eDNA captured on that filter (Takahashi et al., 2023). An alternative and increasingly explored approach is passive sampling, which involves deploying sampling materials (eg: granular adsorbents or membranes) directly in the water for a defined period, allowing eDNA to accumulate through adsorption and/or physical entrapment onto the material (Bessey et al., 2021; Kirtane et al., 2020; Sandré et al., 2026). The idea of passive sampling for aquatic monitoring was first developed for microbial surveillance using Moore swabs in 1948, then used and developed further for the sampling of chemical contaminants, and only applied specifically to eDNA contexts since the 2020s (Godlewska et al., 2021; Huckins et al., 1990; Kirtane et al., 2020; Moore, 1948).

eDNA passive sampling offers several practical advantages that support its continued development. It reduces the need for bulky filtration equipment, allows multiple replicates to be deployed in parallel, and requires less specialized training in the field (Bessey et al., 2021; Jeunen et al., 2022; Kirtane et al., 2020). DNA concentrations can fluctuate rapidly in response to biological activity and hydrological events (Barnes & Turner, 2016; Jensen et al., 2022), causing many species to occur at concentrations near analytical detection limits and increasing the risk of false-negative detections from individual grab samples (Ficetola et al., 2015; Wilcox et al., 2013). Passive samplers therefore have the potential to integrate transient eDNA signals over the deployment period while accumulating low concentrations of eDNA, reducing the likelihood that episodic events or rare taxa are missed by individual grab samples (Bessey et al., 2021; Kirtane et al., 2020; Sandré et al., 2026). However, whether these proposed advantages consistently translate into improved species detection remains unclear (Sandré et al., 2026).

Ideally, a passive sampling approach integrates temporal variation in eDNA availability into a time-weighted average sample (Kirtane et al., 2020; Sandré et al., 2026). For an ideal time-weighted average signal, passive samplers should capture eDNA at a constant rate over time while preserving the captured material by limiting loss until laboratory analysis. Thus, the development and accurate interpretation of passively captured eDNA signals depend on a mechanistic understanding of the kinetics of DNA capture and its subsequent retention on the sampler. If binding sites on the sampler saturate rapidly, early-arriving eDNA will dominate the signal. Conversely, if eDNA captured on the passive sampler is lost over time, more recently captured eDNA will be preferentially retained. In both cases, the recovered eDNA signal reflects a biased integration over time rather than a true time-weighted representation of the underlying eDNA signal. The mechanisms governing eDNA capture and retention on passive sampling materials remain largely unexplored, despite their central role in the functioning of the samplers.

A fundamental challenge in understanding passive eDNA sampling mechanistically is that eDNA is considerably more complex than the dissolved chemical contaminants for which most of the passive sampling theory has been developed (Godlewska et al., 2021). Aquatic eDNA exists in multiple states, including membrane-bound DNA (i.e., DNA encapsulated within cellular or organellar structures), dissolved extracellular DNA, and particle-associated or adsorbed DNA (Mauvisseau et al., 2021; Nagler et al., 2022). These states differ in their persistence, biodiversity information, and transport dynamics and are therefore likely to influence both eDNA capture and retention on passive samplers (Kirtane et al., 2025; Mauvisseau et al., 2021; Nagler et al., 2022). Consequently, the recovered passive sampling signal likely reflects the combined capture of multiple eDNA states instead of a single, uniform process. Despite the likely contribution of multiple eDNA states to passive sampling, dissolved extracellular DNA has been proposed as an important target because adsorption to the sampler surface may protect it from degradation, thereby promoting its long-term retention (Cai et al., 2006; Demaneche et al., 2001; Kirtane et al., 2020). However, eDNA states have not yet been considered or experimentally evaluated in the development of passive samplers.

Deployment time is one of the most important experimental parameters in passive eDNA sampling. Since the first studies of passive eDNA sampling, rapid saturation has been a recurring observation, reflected by both the plateauing of target eDNA accumulation measured by qPCR (Kirtane et al., 2020) and the stabilization of species detection measured by metabarcoding (Bessey et al., 2021), with samplers saturating within minutes to hours of deployment (Bessey et al., 2022; Zhang et al., 2024). A recent meta-analysis further showed that increasing submergence time does not consistently improve species detection across freshwater and marine systems and can even reduce detection due to degradation and biofouling (Sandré et al., 2026). Among the materials tested in freshwater systems, only glass fiber membranes have demonstrated continued eDNA accumulation over time, with uptake reported for up to 72 hours for the Chinese giant salamander (Chen et al., 2022). However, little is known about whether passive eDNA capture kinetics differ among species. Additionally, it remains unknown whether captured eDNA is retained on passive sampling materials throughout the deployment period or continually replaced by newly captured eDNA.

The choice of organisms for addressing these questions should combine strong practical relevance with well-characterized species-specific qPCR assays. Invasive bivalves, including zebra mussels (*Dreissena polymorpha*), quagga mussels (*Dreissena bugensis*), and Asian clams (*Corbicula fluminea*), are well known for their capacity to disrupt aquatic ecosystems and damage critical drinking water infrastructure (Connelly et al., 2007; Karatayev et al., 2015; Sousa et al., 2008; Strayer, 2009). Early detection of these species is essential for effective management of their spread, and research aimed at improving detection methods is therefore urgently needed. As a result, these taxa have been widely targeted in eDNA-based monitoring efforts and have served as model systems for developing and evaluating new sampling approaches (De Ventura et al., 2017; Kirtane et al., 2024; Schabacker et al., 2020; Sepulveda et al., 2019; Suzuki, Houki, et al., 2023). These bivalve species are not only ecologically important, but also share similar functional traits, particularly filter feeding, making them ideal functional replicates for evaluating passive eDNA sampler performance across closely related ecological strategies. Current methods rely exclusively on active eDNA sampling, making invasive bivalves an ideal model organism for evaluating the mechanisms and performance of passive samplers.

Here, we use these three invasive bivalve species and glass fibre membranes as passive sampling material in a controlled mesocosm setting to address three key questions: 1) When do glass fibre samplers saturate with bivalve eDNA, and do these dynamics differ among species? 2) Once captured, how long does bivalve eDNA remain preserved on the samplers? 3) How do accumulation dynamics and retention differ between membrane-bound and dissolved states of eDNA?

## Methods

### Sample collection, mesocosms, and eDNA sources

We collected zebra mussels (*Dreissena polymorpha*) and Asian clams (*Corbicula fluminea*) by hand while snorkelling at a depth of approximately 1 m in Lake Zürich, Switzerland (47.223654, 8.946453) on 3 June 2025. We collected quagga mussels (*Dreissena bugensis*) by hand at a similar depth in Lake Constance, Switzerland (47.644695, 9.212199) on 23 June 2025. We maintained all bivalves at the Aquatikum facility of the Swiss Federal Institute of Aquatic Science and Technology (Eawag) in aerated holding tanks under contained conditions that prevented accidental release into the environment. We replaced 50% of the holding tank water every two weeks and fed the bivalves a culture of the green alga *Chlamydomonas reinhardtii* at the same interval. Water temperature remained constant at approximately 15.5 °C throughout the experiments.

We conducted experiments in nine mesocosms (35 × 19 × 15 cm; L × W × H), each containing 6.5 L of water and one of three invasive bivalve species: quagga mussels, zebra mussels, or Asian clams. Each mesocosm contained ten individuals of a single species, with three replicate mesocosms per species. We recorded the total biomass of bivalves and the physicochemical characteristics of the mesocosm water (Tables S1 and S2). Throughout the experiment, we continuously aerated each mesocosm using air bubblers to maintain mixing throughout the water column. We did not replace the water or feed the bivalves during the experimental period.

To account for multiple states, we used dissolved DNA spikes to isolate passive capture dynamics of the dissolved state in addition to the eDNA released by the bivalves. Specifically, we spiked mesocosms with extracted DNA from chum salmon (*Oncorhynchus keta*), chicken (*Gallus gallus*), and mouse (*Mus musculus*). We obtained sheared salmon sperm DNA (10 mg mL⁻¹) from a commercial supplier (Invitrogen; catalog no. AM9680). We extracted mouse DNA from cultured cells and chicken DNA from store-bought chicken following Kirtane et al., 2023. Prior to spiking, we quantified all DNA extracts using qPCR (Table S4). We then added 1 mL of the respective DNA extract to each mesocosm across all experiments, pairing salmon with quagga mussels, mouse with Asian clams, and chicken with zebra mussels. This yielded final concentrations in the tank water of ∼2,690, ∼43,623, and ∼905,620 DNA copies/mL of mesocosm water for salmon, mouse, and chicken, respectively (Table S4).

### Passive samplers

We used 25 mm diameter glass fibre membranes (1 µm nominal pore size; Whatman, catalog no. WHA1821025) as passive eDNA samplers, as this material exhibits a steady increase in eDNA capture over time in freshwater systems and outperformed active sampling in field conditions (Chen et al., 2022; Qian et al., 2026). We deployed the glass fibre membranes in specifically designed 3D-printed housings and suspended them within mesocosm tanks using modified 3-inch 3D-printed hooks (Figure S1; Schadewell et al., 2026). In all experiments, we positioned the samplers in the mid-water column, approximately 6 cm below the water surface. All membrane housings were sterilized with 10% bleach before assembly and deployment. Figure S1 shows photographs of the experimental setup.

### Experiment 1: eDNA temporal dynamics

To quantify the temporal dynamics of eDNA on passive samplers, we established nine mesocosms (three per treatment), each containing nine passive sampling membranes arranged across three triplicate housings (three membranes per housing) (Figure 1A, S1). Treatments consisted of 1) 10 quagga mussels with a salmon DNA spike, 2) 10 zebra mussels with a chicken DNA spike, and 3) 10 Asian clams with a mouse DNA spike. Mussels and clams were maintained in the mesocosms for two weeks before the experiment to allow eDNA concentrations to reach a steady state (Sansom & Sassoubre, 2017). Each mesocosm was continuously aerated using a bubbler to oxygenate and mix the water. Immediately before sampler deployment, we added 1 mL of dissolved DNA from the corresponding spike species to each mesocosm.

**Figure 1:**
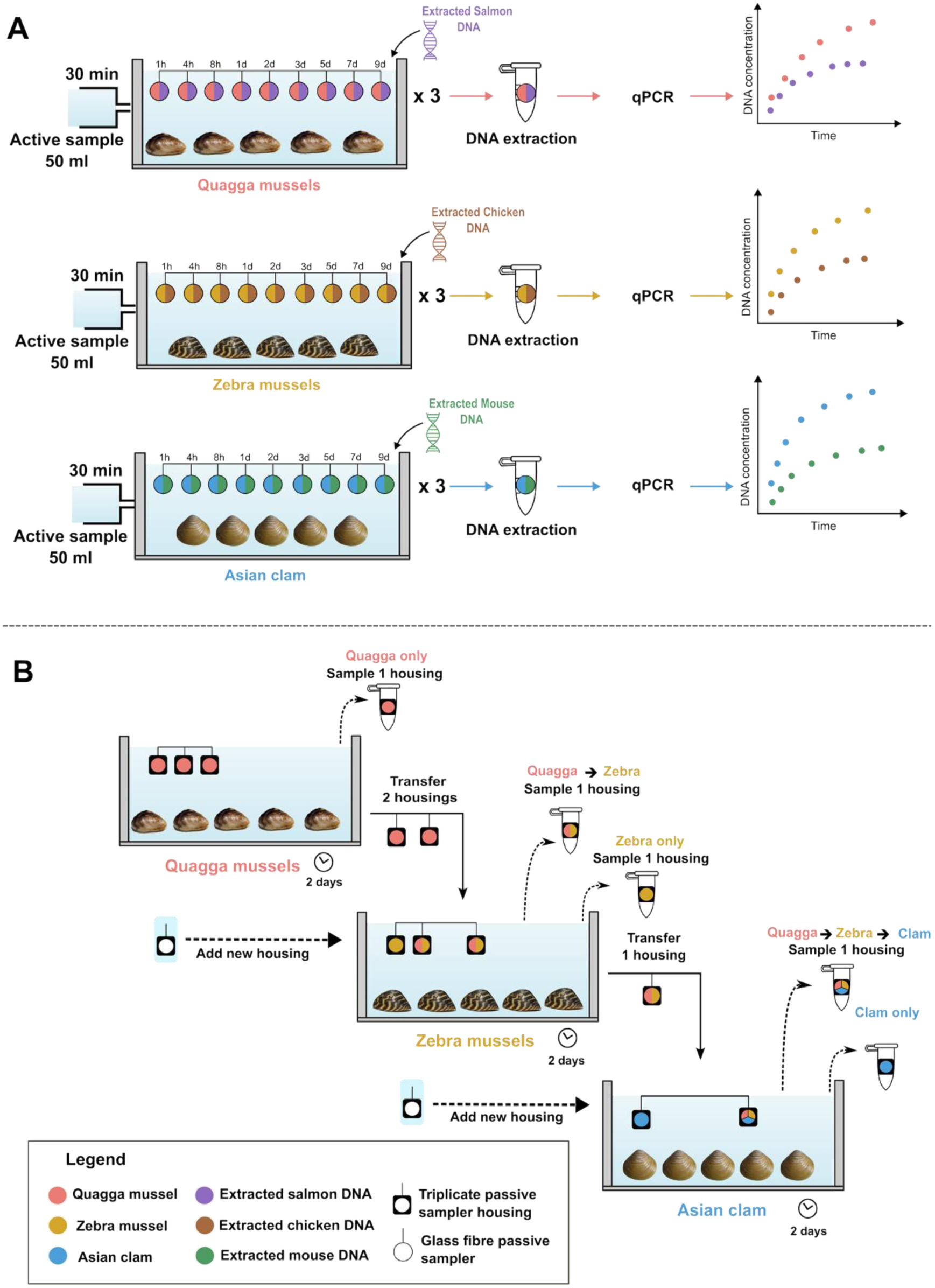
Experimental design of the passive eDNA sampling experiments. (A) Illustration of the eDNA temporal dynamics experiment. (B) Illustration of the eDNA transfer experiment. Different colors represent DNA originating from different species and illustrate the expected composition of eDNA detected by passive samplers throughout the experiments. Triplicate passive sampler housings hold three glass fibre membranes (see Figure S1).

We collected the deployed membranes from each mesocosm at 1 h, 4 h, 8 h, 1 d, 2 d, 3 d, 5 d, 7 d, and 9 d time points. During retrieval, we removed one membrane from its triplicate housing, stored in 600 µL of Longmire lysis buffer (100 mM Tris pH 8.0, 0.5 mM EDTA, 0.5% SDS, 200 mM NaCl), at −20 °C until extraction. We maintained a negative control mesocosm containing no mussels and no added DNA alongside the experimental tanks. In this control tank, we deployed one triplicate housing with three membranes, and sampled them at 1 d, 3 d, and 9 d timepoints. We processed control samples in parallel with experimental samples to assess potential contamination.

### Experiment 2: eDNA retention test

To investigate the persistence and replacement dynamics of eDNA captured on passive samplers, we conducted a sequential transfer mesocosm experiment (Figure 1B). We reused the same nine mesocosms following completion of the temporal dynamics experiment. We treated each triplicate housing (containing three membranes) as the experimental unit and transferred or sampled all three membranes in the housing as intact units. We exposed the triplicate housings for fixed 2-day intervals and sequentially transferred them between mesocosms containing different species.

In the initial phase, we deployed three triplicate housings in each of the quagga mussel mesocosms. After 2 days, we collected one triplicate housing from each mesocosm and transferred the remaining housings to zebra mussel mesocosms. At the time of transfer, we deployed an additional set of fresh triplicate housings to enable comparison between transferred and newly deployed samplers. After a further 2-day deployment in zebra mussel mesocosms, we collected one transferred and the newly deployed housings from each tank. We then transferred the one remaining housing to Asian clam mesocosms. At this stage, we again introduced a new triplicate housing. Following a final 2-day deployment, we collected all deployed housings from the Asian clam mesocosms. Across the experiment, this design generated five passive sampler histories: (i) quagga-only, (ii) zebra-only, (iii) Clam-only, (iv) quagga→zebra transfer, and (v) quagga→zebra→Clam transfer. In total, we collected 24 triplicate sampler housings (72 individual membranes). After sampling, we transferred each membrane into 2 mL tubes containing 600 µL of Longmire’s lysis buffer and stored them at −20 °C until extraction.

### Experiment 3: active vs passive capture

To contextualize eDNA recovery by passive samplers, a 50 mL active sample was collected from each mesocosm tank 30 min after the addition of dissolved DNA during the temporal dynamics experiment (experiment 1). The water sample was then filtered through the same glass fibre membrane type using a Swinnex filter holder and a sterile syringe. The syringes and filter holders were sterilized with 10% bleach prior to the experiment. The membranes from active samples were processed using the same DNA extraction and qPCR workflow as the passive sampler membranes. Since passive sampling membranes were retrieved at multiple time points throughout Experiment 1, membranes collected at time point of 9 days were used for the mussel targets (Quagga mussel, Zebra mussel, and Asian clam), whereas membranes collected at the 8-h time point were used for the dissolved DNA targets (salmon, chicken, and mouse). These time points correspond to the highest concentration of eDNA for each target group in Experiment 1 captured by the passive sampling membranes (Figure 2).

**Figure 2.**
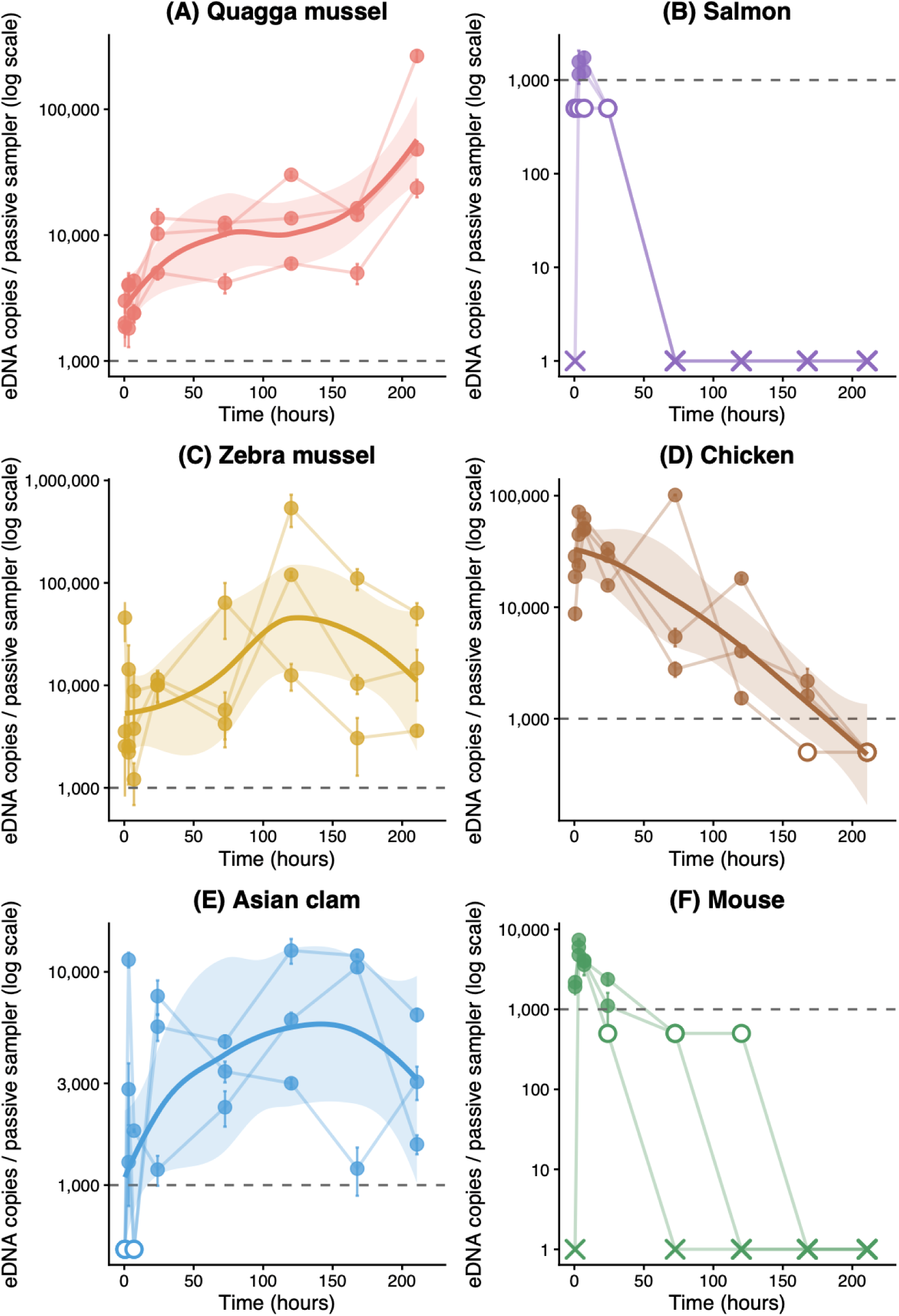
eDNA concentration (copies per passive sampler membrane) over time for bivalve-derived eDNA (left column: (A) Quagga mussel, (C) Zebra mussel, and (E) Asian clam) and dissolved eDNA (right column: (B) Salmon, (D) Chicken, and (F) Mouse). Points represent the mean concentration for each mesocosm at each sampling time, with error bars indicating the standard error among qPCR replicates. Lines connect repeated measurements from the same mesocosm. Quantifiable values are shown as filled circles; values below the limit of quantification are plotted as open circles at 500 copies/sampler for visualization, while non-detections are indicated by crosses. The dashed horizontal line indicates the limit of quantification (1000 copies per sampler).

### DNA extraction

We thawed all collected membranes and added 15 µL of proteinase K (20 mg mL⁻¹) to each tube. We incubated samples for 3 h at 56 °C with shaking at 300 rpm. Following incubation, we transferred 550 µL of lysate to a 2 mL 96-well plate containing 415 µL of hybridization buffer (for 500 mL: 1 g DTT, 72.5 g NaCl, 125 g PEG 8000, 500 µL 0.5 M EDTA, brought to 500 mL with molecular-grade water) and 20 µL of magnetic beads (SeraMag SpeedBeads, carboxylate-modified magnetic particles, hydrophobic; Cytiva, catalog no. 65152105050250). DNA extraction was performed using a KingFisher Apex system (Thermo Fisher Scientific, Waltham, USA). The solution and beads were mixed for 10 min (slow mode) to facilitate DNA binding. Beads were then collected using the magnetic head and held above the wells for 1 min to allow excess liquid to drain. This was followed by two sequential washes in 800 µL of 80% ethanol, each with 1 min mixing (slow mode) and subsequent bead collection. After washing, beads were air-dried for 10 min to remove residual ethanol. DNA was eluted in 100 µL of TE buffer preheated to 70 °C, with 1 min mixing (slow mode) prior to bead collection. Three extraction controls were included and interspersed across the plate to monitor potential contamination during the extraction process.

### Quantitative PCR

We quantified all target DNA concentrations from all membranes using previously published species-specific probe-based qPCR assays (Table 1). We performed reactions in 384-well plates on a Roche LightCycler 480 in a total volume of 10 µL, consisting of 5 µL of 2× TaqMan Gene Expression Master Mix (Applied Biosystems), 0.05 µL of each primer (500 nM final concentration), 0.025 µL of probe (250 nM final concentration), and 3.875 µL of nuclease-free water. We prepared qPCR plates using a mosquito® LV liquid handler (SPT Labtech, Melbourn, UK). For all assays except the zebra mussel assay, we used thermal cycling conditions consisting of an initial enzyme activation at 95 °C for 10 min, followed by 40 cycles of denaturation at 95 °C for 15 s and annealing/extension at 60 °C for 45 s. For the zebra mussel assay, we applied modified cycling conditions with an initial denaturation at 95 °C for 10 min, followed by 55 cycles of denaturation at 95 °C for 1 min, annealing at 55 °C for 1 min, and extension at 72 °C for 30 s. We included a gBlock six-point standard curve (10⁶–10 copies per reaction) in each qPCR run (Table S3). We analyzed all samples in triplicate, all standards in six replicates, and included at least three no-template controls per plate to monitor potential contamination. We determined Cq values using a fixed fluorescence threshold and converted them to copy numbers using standard curve regression. We quantified samples on each plate using the corresponding standard curve generated on that plate.

**Table 1:**
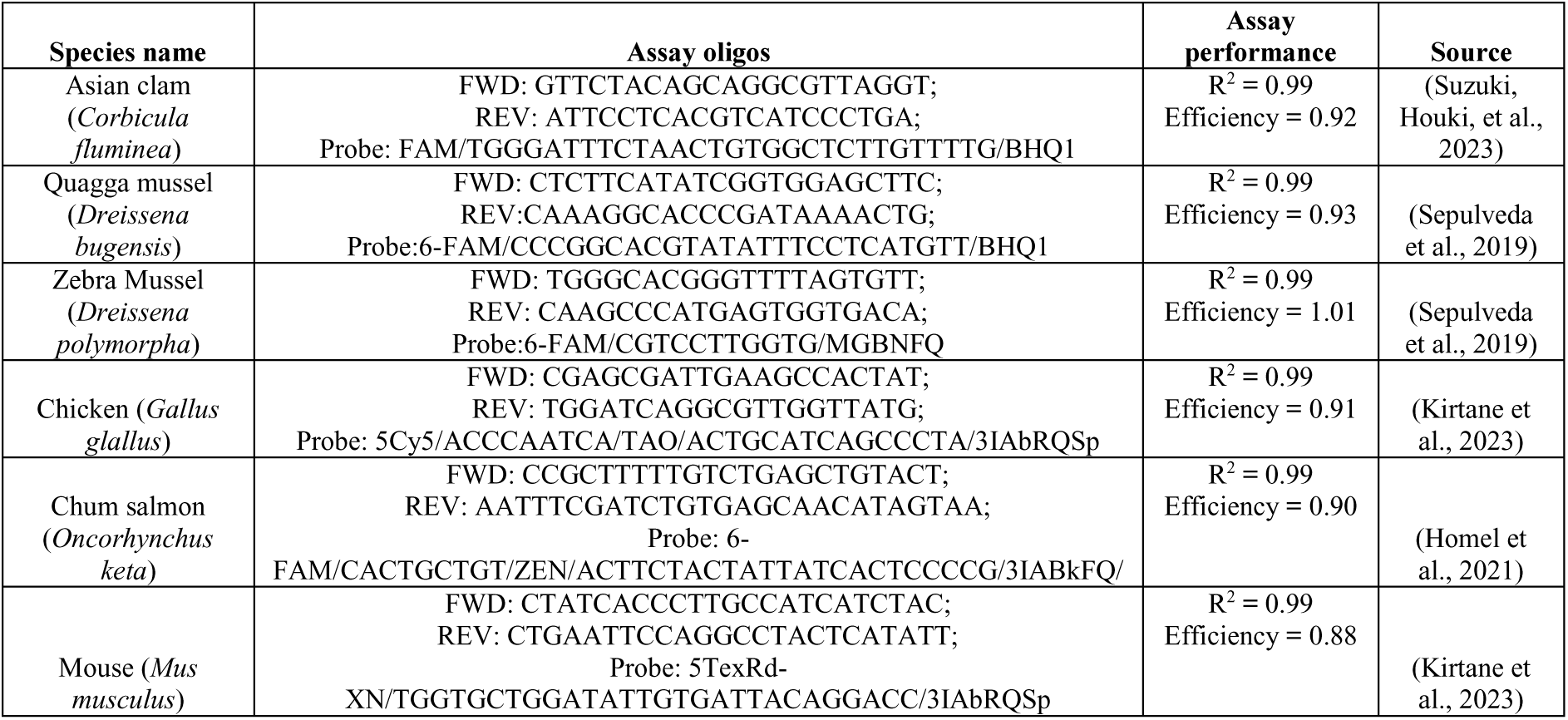
qPCR assays for all targets used in this study.

### Data analyses

#### Quantification of eDNA copies

We determined Cq values using a fixed fluorescence threshold and converted them to copy numbers using the standard curve regression on the same qPCR plate. Average standard curve performance statistics for all qPCR assays are provided in Table 1. We defined the assay-specific limit of quantification (LOQ) as the lowest standard curve concentration with a coefficient of variation (CV) below 0.35 (Klymus et al., 2020). Samples were considered non-detects when fewer than three technical (qPCR) replicates amplified. Samples in which all three technical replicates amplified, but the estimated concentration was below the LOQ, were classified as positive detections below the LOQ. These samples were retained in all figures using their estimated concentrations, but for statistical analyses were assigned a value equal to half the assay-specific LOQ. All qPCR assays had average efficiencies between 0.88 and 0.101, R^2^ between 0.98 and 0.99 (Table 1), and a LOQ of 10 copies/qPCR reaction or 1000 copies/membrane. None of the experimental, extraction, or qPCR negative controls showed amplification in all three technical replicates or amplification above the LOQ, indicating a low probability of contamination.

#### Experiment 1: temporal dynamics

Temporal trends in eDNA concentration were assessed using linear models with time as a continuous predictor. Analyses were performed separately for each species using log₁₀-transformed concentrations. Models were fitted to the full dataset and separately to early (≤24 h) and late (>24 h) sampling periods to assess potential phase-dependent dynamics.

#### Experiment 2: sequential transfer experiment

For the sequential transfer experiment, differences in eDNA concentrations among transfer histories, defined as combinations of species exposure and transfer sequence, were assessed separately for each target bivalve species. Treatment histories included quagga-only, zebra-only, clam-only, quagga→zebra, and quagga→zebra→clam. Samples with fewer than three detected technical replicates were treated as non-detections. Differences in eDNA concentrations among treatment histories were assessed using Kruskal–Wallis tests followed by Dunn’s post hoc comparisons with Benjamini–Hochberg correction. Where only two treatment histories were available, differences were assessed using the Wilcoxon rank-sum test.

#### Experiment 3: active vs passive capture

To compare active and passive eDNA sampling, qPCR replicates were averaged for each analyzed membrane. For passive samplers, concentrations from Experiment 1 were used as described above. Concentrations were log₁₀-transformed prior to analysis. Differences between active and passive sampling were assessed separately for each target using Welch’s two-sample *t*-tests. All statistical analyses were conducted in R version 4.4.2 (R Core Team, 2024).

## Results

### Experiment 1: eDNA accumulation dynamics

#### Bivalve eDNA accumulation is species-specific

All three bivalve species showed a significant positive relationship between eDNA concentration and time (Table 2). When the data were split into early (≤24 h) and late (>24 h) capture phases, patterns differed among species. Passive sampling of quagga mussel eDNA showed a significant positive relationship with time in both phases. Asian clam eDNA showed a significant positive relationship during the first 24 hours, but not during the later phase. In contrast, zebra mussel eDNA showed no significant relationship with time in either phase (Table 2). After 9 days of passive deployment, Asian clam eDNA reached a mean concentration of approximately 3,100 copies per membrane (range: 1,540–6,235), compared with approximately 13,900 copies (3,690–52,300) for zebra mussels and 67,200 copies (19,400– 232,300) for quagga mussels. Thus, passive samplers accumulated approximately 4.5-fold more zebra mussel eDNA and over 20-fold more quagga mussel eDNA than Asian clam eDNA after 9 days of deployment, although the inter-membrane variability was consequent.

**Table 2.** Linear model results for full, early (≤24 h), and late (>24 h) sampling phases, describing the relationship between eDNA concentration and time for three mussel species (Quagga mussel, Zebra mussel, Asian clam) and spiked extracted DNA from chicken. Linear models for spiked DNA from mouse and salmon were not run due to too few data points.

| Species | Phase | Slope (Estimate) | Standard Error | t-value | p-value |
| --- | --- | --- | --- | --- | --- |
| Asian clam | Full | 0.00250 | 0.000687 | 3.64 | $5.26 \times 10^{-4}$ |
| Asian clam | Early | 0.0250 | 0.00837 | 2.98 | $5.41 \times 10^{-3}$ |
| Asian clam | Late | -0.000305 | 0.00112 | -0.27 | 0.787 |
| Quagga mussel | Full | 0.00528 | 0.000468 | 11.3 | $4.87 \times 10^{-19}$ |
| Quagga mussel | Early | 0.0256 | 0.00372 | 6.87 | $1.78 \times 10^{-8}$ |
| Quagga mussel | Late | 0.00556 | 0.00110 | 5.06 | $7.26 \times 10^{-6}$ |
| Zebra mussel | Full | 0.00343 | 0.00164 | 2.09 | $4.00 \times 10^{-2}$ |
| Zebra mussel | Early | 0.0158 | 0.00974 | 1.62 | 0.114 |
| Zebra mussel | Late | 0.00725 | 0.00447 | 1.62 | 0.114 |
| Chicken (DNA) | Full | -0.00858 | 0.000601 | -14.3 | $1.93 \times 10^{-22}$ |
| Chicken (DNA) | Early | -0.00122 | 0.00496 | -0.25 | 0.808 |
| Chicken (DNA) | Late | -0.0100 | 0.00154 | -6.50 | $1.93 \times 10^{-7}$ |

#### Dissolved eDNA degrades rapidly after introduction

In contrast to mussel-derived eDNA, dissolved DNA from chicken, mouse, and salmon showed an initial increase during the first eight hours, after which concentrations declined. Statistical analysis could only be conducted for the chicken spike, as mouse and salmon concentrations rapidly fell below the limit of quantification. Nevertheless, both mouse and salmon DNA were consistently detected during the early sampling period but were largely absent after 24 h (Figure 2). Chicken DNA, which had higher starting concentrations (Table S4), exhibited a strong decline over time (estimate = −0.0086, *p* < 0.001), showing a significant negative slope. When the data were split into early (≤24 h) and late (>24 h) phases, no significant relationship with time was observed during the early phase. However, during the late phase, chicken DNA showed a significant negative relationship with time, indicating continued degradation (Table 2). Furthermore, the concentration of chicken DNA in the mesocosm water corresponded to approximately 9 × 10⁵ copies/mL (Table S4). Based on an estimated pore water volume of 200–300 µL retained within each passive sampler, this corresponds to an expected range of ∼1.8 × 10⁵ to 2.7 × 10⁵ copies per sampler. The observed maximum concentrations (∼1 × 10⁵ copies/sampler) were of the same order of magnitude (Figure 2D).

### Experiment 2: Loss of accumulated eDNA following sequential transfer

The eDNA concentrations of quagga mussels differed significantly across transfer histories (Kruskal–Wallis test, χ² = 40.99, df = 2, p = 1.26 × 10⁻⁹). Following transfer to zebra mussel mesocosms, the concentrations of quagga mussel eDNA in passive sampler membranes declined, with many replicates falling below the limit of quantification (Figure 3A; Dunn test, p_adj = 7.41 × 10⁻⁵). A further decrease in quagga mussel eDNA concentrations was observed after subsequent transfer to Asian clam mesocosms for an additional two days (Figure 3A; Dunn test, p_adj = 2.00 × 10⁻⁹). However, this reduction was not statistically significant relative to the preceding zebra mussel mesocosm exposure (Dunn test, p_adj = 0.062) because by the final transfer, most samples were already either not detected or below the limit of quantification, which limited our ability to statistically resolve any additional decline (Figure 3A).

**Figure 3:**
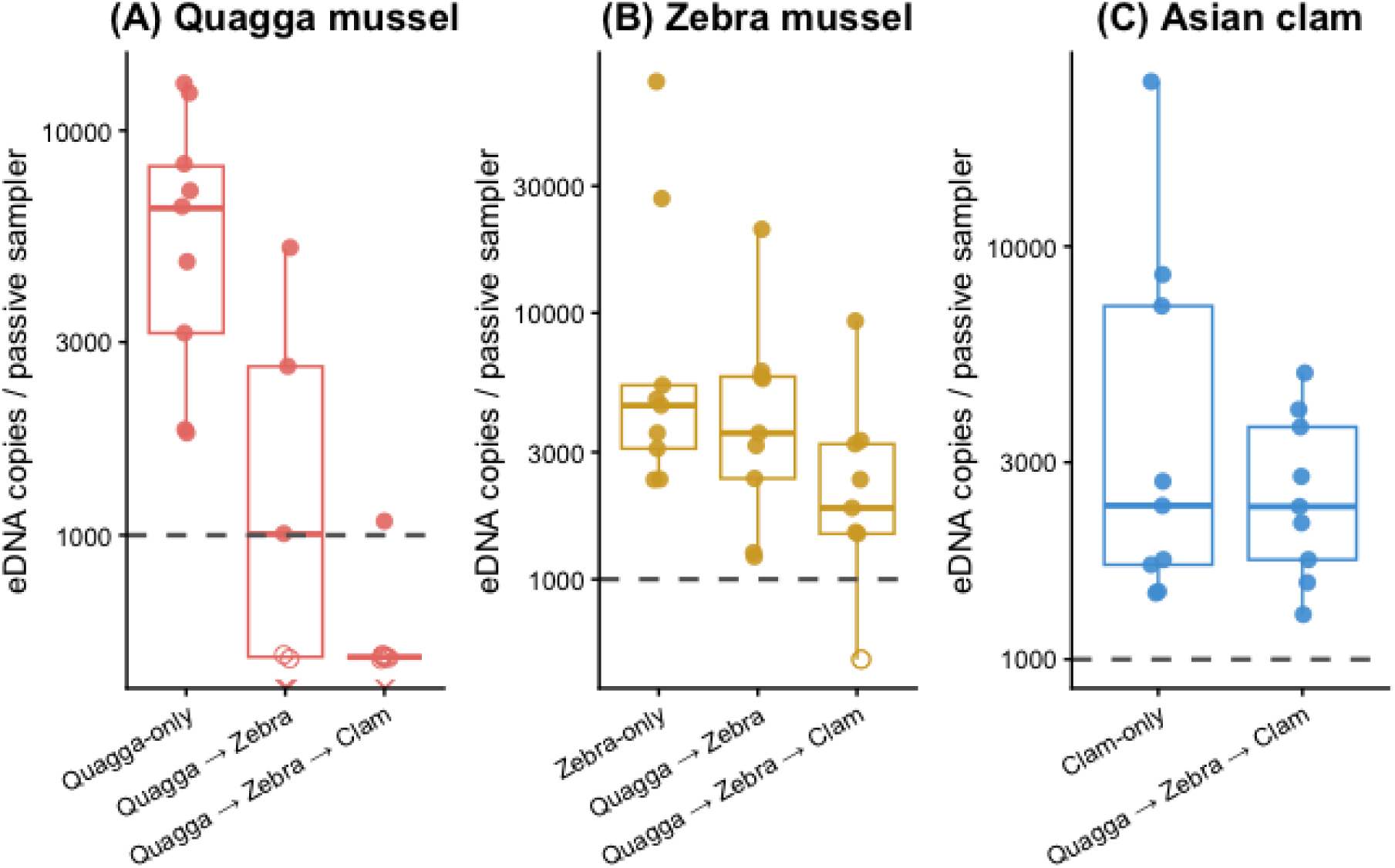
eDNA concentrations (copies per passive sampler membrane) across treatment histories for (A) Quagga mussel, (B) Zebra mussel, and (C) Asian clam. Points represent average eDNA concentration from each membrane (n = 9 per treatment), and boxplots summarise their distribution. Quantifiable values are shown as filled circles; values below the limit of quantification are plotted as open circles at 500 copies/sampler for visualization, while non-detections are indicated by crosses. The dashed horizontal line indicates the limit of quantification (1000 copies/sampler).

The eDNA concentrations of zebra mussels differed significantly across transfer histories (Kruskal–Wallis test, χ² = 14.71, df = 2, p = 6.38 × 10⁻⁴). After transfer to the Asian clam mesocosms, zebra mussel eDNA concentrations declined significantly after two days (Figure 3B), with concentrations in the quagga → zebra → clam transfer history lower than both the preceding zebra-only (Dunn test, padj = 0.0005) and the quagga → zebra histories (Dunn test, padj = 0.017). Following exposure to zebra mussel mesocosms, eDNA concentrations of zebra mussels were comparable between the zebra-only and quagga → zebra histories (Dunn test, p_adj = 0.215), indicating no detectable difference in the accumulation of zebra mussel eDNA irrespective of prior exposure to quagga mussels (Figure 3B). Similarly, the eDNA concentrations of the Asian clam did not differ significantly between the clam-only and the quagga → zebra → clam transfer histories (Figure 3C; Wilcoxon rank-sum test, W = 402, p = 0.522).

### Experiment 3: active vs passive capture

Passive samples yielded higher quagga mussel eDNA concentrations than active sampling; however, this difference was not statistically significant (Figure 4A; Welch’s two-sample *t*-test, *p* = 0.30). In contrast, passive samples recovered lower eDNA concentrations than active sampling for salmon (Figure 4B) and zebra mussels (Figure 4C; *p* = 0.33), although neither difference was statistically significant. For salmon, all active sample concentrations were consistently below the assay LOQ, while some replicates of passive sampling were above the LOQ (Figure 4B). Active sampling recovered significantly higher eDNA concentrations than passive sampling for chicken (Figure 4D; *p* = 0.001), Asian clam (Figure 4E; *p* = 0.042), and spiked mouse DNA (Figure 4F; *p* = 0.010). Overall, differences between active and passive sampling were target-dependent, with active sampling recovering significantly higher eDNA concentrations for Asian clam and two of the three dissolved DNA spikes, whereas no significant differences were observed for the remaining targets.

**Figure 4:**
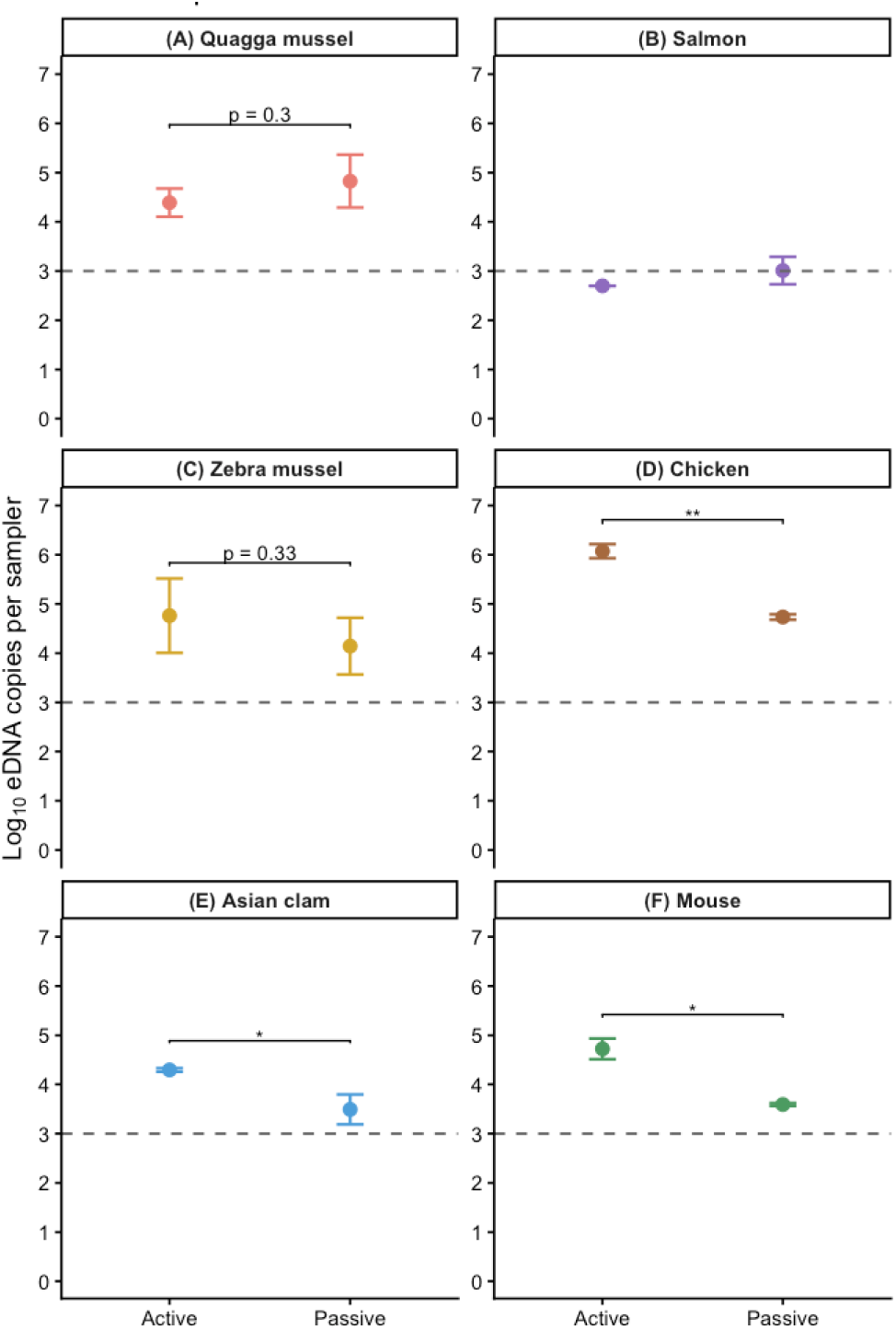
Comparison of active (50 mL water sample) and passive eDNA sampling for (A) Quagga mussel, (B) Salmon, (C) Zebra mussel, (D) Chicken, (E) Asian clam (Corbicula), and (F) Mouse. Passive samplers were deployed for 9 days in the invasive bivalve mesocosms and for 1 h in the dissolved eDNA spike experiments. Points and error bars represent the mean ± standard deviation of biological replicates (n = 3). The dashed horizontal line indicates the limit of quantification (1000 copies per sampler). Asterisks indicate significant differences between active and passive sampling based on Welch’s two-sample t-tests (p < 0.05)

## Discussion

Here, we used glass fibre membranes in a controlled mesocosm experiment to investigate the accumulation and retention dynamics of passive eDNA sampling. Specifically, we asked (1) when passive samplers saturate with eDNA and whether these dynamics differ among invasive bivalve species, (2) how long captured eDNA remains associated with the membranes following transfer between mesocosms, and (3) how accumulation dynamics differ between membrane-bound and dissolved eDNA. We found that eDNA accumulation dynamics depended strongly on both species and eDNA state. Passive samplers rapidly accumulated eDNA from the living bivalves before reaching species-specific plateaus, whereas dissolved eDNA declined over the same deployment period. Transfer experiments showed that previously captured eDNA was rapidly replaced by the local signal, indicating that glass fibre membranes might primarily reflect recent exposure.

### eDNA accumulation dynamics are state-specific and species-specific

The spiked dissolved DNA from chicken declined in concentration throughout the experiment, consistent with previous studies suggesting that dissolved eDNA is relatively vulnerable to degradation unless protected through adsorption to particles or surfaces (Mauvisseau et al., 2021; Nagler et al., 2018). Our experiment suggests that adsorption to the glass fibre membranes, if it occurred, was insufficient to prevent losses. This contrasts with previous work demonstrating the adsorption of dissolved DNA onto glass fibre surfaces (Schadewell et al., 2026; Suzuki, Kawamura, et al., 2023), though Schadewell et al. also observed a sharp decline in the captured DNA concentration within the first 24 hours. For the dissolved DNA spikes from salmon and mouse, a consistent pattern of early detection followed by a rapid decline to below the limit of quantification, suggesting rapid loss of dissolved eDNA. These findings should be interpreted with caution given the lower initial target concentrations of mouse and salmon spikes, and highlight the consideration for higher dissolved DNA concentrations in future studies.

For bivalves, the eDNA accumulation did not continue throughout the deployment for all species. Accumulation ranged from little change after deployment (zebra mussel), to an early plateau (Asian clam), to a continuous increase throughout the 9-day deployment (quagga mussel). These contrasting accumulation dynamics suggest that species differ in the way eDNA is released and subsequently captured by passive samplers. High variability in the concentration of bivalve eDNA copy number was observed among replicate samplers and mesocosms, sometimes over an order of magnitude. This could be explained by the variability in eDNA shedding rates and in the physical form/state of eDNA (e.g. dissolved DNA, intact cells, damaged cells, or cellular aggregates), which may impact their accumulation on glass fibre membranes (Mauvisseau et al., 2021; Power et al., 2023).

Body size influences eDNA shedding rates in some species, although the direction and magnitude of this relationship appear to vary among taxa (Maruyama et al., 2014; Takeuchi et al., 2019). This relationship may not apply to bivalves because much of their total mass comprises the shell rather than metabolically active tissue. Consistent with this, Asian clams were among the heaviest individuals used in this study but accumulated the lowest eDNA concentrations, whereas zebra mussels, despite being the lightest, accumulated comparatively high eDNA concentrations. Species-specific differences in filtration rates may also have contributed to the contrasting eDNA concentrations observed among species. Zebra and quagga mussels tend to have higher filtration rates than Asian clams (Marescaux et al., 2016;

Silverman et al., 1995), consistent with the comparatively lower eDNA concentration of the clams captured in the passive samplers. Furthermore, bivalve species differ in the production and characteristics of biodeposits, including mucus-rich feces and pseudofeces, which transform fine suspended particles into larger aggregates (Atkinson et al., 2011; Murphy et al., 2019). If eDNA is incorporated into these biodeposits, it could change their persistence, settling behavior, and capture by passive samplers compared to dissolved DNA or individual cells (Graf & Rosenberg, 1997; Li et al., 2008). The intermittent release and passive capture of biodeposits could also contribute to the high temporal and among-membrane variability observed in this study. Future studies should determine how differences in filtration behaviour and biodeposition influence eDNA shedding and passive sampler capture, as these processes may underpin the species-specific accumulation dynamics observed here.

### eDNA does not persist on glass fibre passive samplers

Sequential transfer experiments showed that the concentration of previously accumulated eDNA declined following transfers between mesocosms. Following transfer, eDNA concentrations of the new target on transferred samplers were similar to those measured on freshly deployed samplers in the new mesocosm. Together, these observations indicate that passively captured eDNA on glass fibre membranes is continually lost and replaced. Consequently, glass fibre passive samplers may not provide a true time-weighted average of past conditions but instead become dominated by more recent exposures.

The observed loss in eDNA concentration following sequential transfer could have resulted from degradation, desorption, or other removal processes. The disappearance of previously accumulated eDNA within 24–48 h is consistent with reported persistence times of bivalve eDNA (Ruiz-Ramos et al., 2024; Sansom & Sassoubre, 2017) and of spiked dissolved DNA observed in Experiment 1 (Figure 2). Since eDNA capture continued while previously captured eDNA was simultaneously lost, the observed plateau for zebra mussels and Asian clams is unlikely to reflect saturation of available binding sites on the passive sampler. Instead, the plateau may be consistent with a dynamic equilibrium in which ongoing capture is balanced by concurrent loss processes, analogous to the steady-state eDNA concentrations commonly observed in the water column (Sansom & Sassoubre, 2017). This suggests that eDNA concentrations on glass fibre passive samplers are regulated by dynamic turnover rather than continuous accumulation.

The observed capture dynamics also provide insight into which eDNA states contribute most to the passive sampler signal. For instance, the dissolved DNA experiment showed little evidence of accumulation, as the highest concentrations recovered from passive samplers were comparable to those expected from the pore water retained within the glass fibre membrane. This suggests that dissolved eDNA is unlikely to be the dominant contributor to long-term signal retention. In contrast, the mesh-like structure of glass fibre membranes is likely conducive to the capture of membrane-bound eDNA (e.g. intact cells, organelles, and cell aggregates) and other particulate material, including biofilm-associated and adsorbed eDNA (Kirtane et al., 2023; Power et al., 2023; Qian et al., 2026). The substantial variability observed among replicate samplers (Figures 2, 3) is also consistent with the stochastic capture of discrete membrane-bound/particulate eDNA instead of the uniform accumulation of dissolved DNA.

### Limitations

This study has some limitations that should be considered when interpreting the results and applying them to more general contexts. First, experiments were conducted under controlled lentic mesocosm conditions, and the capture dynamics observed here may differ under more complex field conditions where hydrology, biofouling, suspended particles, and environmental variability influence eDNA states, transport, and persistence (Barnes et al., 2014; Mauvisseau et al., 2021; URycki et al., 2024). Recent studies have shown that glass fibre passive sampling membranes may perform more effectively in higher flow lotic conditions (Qian et al., 2026). Second, only glass fibre membranes were evaluated in this study. Comparing alternative passive sampling materials with different physical and chemical properties represents an important next step towards optimizing passive eDNA sampling for different monitoring applications (Bessey et al., 2022; Chen et al., 2022). Third, although the results suggest that membrane-bound or particulate eDNA likely dominates the passive sampler signal, the individual contributions of dissolved, membrane-bound, and adsorbed states of bivalve eDNA could not be quantified directly. Likewise, the relative importance of degradation, desorption, and other removal processes contributing to eDNA loss could not be distinguished. Finally, the experiments focused exclusively on invasive freshwater bivalves. Although capture dynamics may differ among organisms with different eDNA shedding characteristics (Andruszkiewicz Allan et al., 2021), the experimental framework developed here provides a basis for testing passive eDNA capture across a broader range of taxa and environmental settings.

### Outlook

There is unlikely to be a single passive sampling strategy that is optimal for all monitoring objectives. Rather, the choice of sampling approach should be guided by the ecological question being addressed and the extent to which eDNA capture mechanisms are understood. For applications such as the early detection of invasive bivalves, the current understanding of passive eDNA accumulation remains insufficient for reliable interpretation of detection probabilities or quantitative comparisons among sites. In these cases, active sampling methods, such as grab samples or tow-net sampling, may be preferred as eDNA concentrations within a fixed volume of water can be interpreted and compared more consistently across samples (Kirtane et al., 2024; Miller et al., 2024; Schabacker et al., 2020; Sepulveda et al., 2019). Furthermore, advances in automated active sampling technologies provide an alternative approach for integrating temporal variation in eDNA signals (Yamahara et al., 2025).

Consideration of eDNA states will be central to the future development of passive eDNA samplers, particularly with respect to the mechanisms governing eDNA retention. For instance, extracellular/dissolved eDNA can be preserved for extended periods through adsorption onto particle surfaces (Cai et al., 2006; Demaneche et al., 2001). Consequently, an ideal passive sampler relying on this mechanism could adsorb the dissolved eDNA at a constant rate while preserving the newly adsorbed DNA until analysis. Knowledge of the capture mechanisms would also guide the choice of appropriate DNA extraction protocols for effective recovery (Kirtane et al., 2020, 2023; Müller et al., 2024). Achieving this remains challenging because multiple factors like material surface properties, ionic composition, dissolved organic matter, and other environmental factors all influence DNA adsorption mechanisms (Kirtane et al., 2020; Müller et al., 2024). Potential solutions can be drawn from aquatic pollutant passive sampling, where environmental effects on analyte uptake are well established (Godlewska et al., 2021). These effects are accounted for through extensive calibration, mechanistic uptake models, and performance reference compounds that estimate in situ sampling rates (Godlewska et al., 2021). Adapting similar approaches to eDNA passive sampling will likely lead to more reliable and interpretable eDNA data.

This study demonstrates that understanding the mechanisms governing passive eDNA sampling is essential for developing passive samplers into practical tools for routine eDNA-based biodiversity monitoring. We show that glass fibre membranes can effectively capture eDNA during short deployment periods; however, extending deployment times will likely require the development or identification of alternative materials with improved capture capacity and retention properties. Future studies should therefore evaluate candidate materials for saturation dynamics, preservation capabilities, and reproducibility in laboratory and field conditions. Lastly, the optimal sampler design and deployment strategy will likely depend on whether the objective is to integrate eDNA signal over time, maximize detection sensitivity, or study temporal changes in eDNA concentrations.

### Artificial Intelligence disclosure

Generative artificial intelligence (AI) tools (ChatGPT, OpenAI) were used during manuscript preparation to assist with language editing and debugging code (R Wizard GPT). All analyses, statistical tests, results, interpretations, and conclusions were designed, verified, and approved by the authors. The authors take full responsibility for the accuracy and content of the manuscript.

## Acknowledgements

We thank Dr. Alex Hall for support and feedback. We thank Dr. Yvonne Schadewell for sharing the 3D-printed membrane housing designs. We thank Pascal Bucher for printing 3D-printing the passive sampler housings. Pascal Bucher and Kirsten Klappert helped in maintaining the bivalve aquariums. We thank Thomas Müller and Linus Hofstetter for collecting quagga mussels. Data produced and analyzed in this paper were generated in collaboration with the Genetic Diversity Centre (GDC), ETH Zurich.

## Supporting information

**Table S1:** Combined mass of ten individuals in each experimental mesocosm at the beginning of the experiments.

| Species | Mesocosm replicate number | Mass (g) |
| --- | --- | --- |
| Quagga mussels | 1 | 25.8 |
|  | 2 | 27.8 |
|  | 3 | 33.0 |
| Zebra mussels | 1 | 15.4 |
|  | 2 | 11.4 |
|  | 3 | 8.4 |
| Asian clam | 1 | 32.0 |
|  | 2 | 30.0 |
|  | 3 | 32.4 |

**Table S2:**
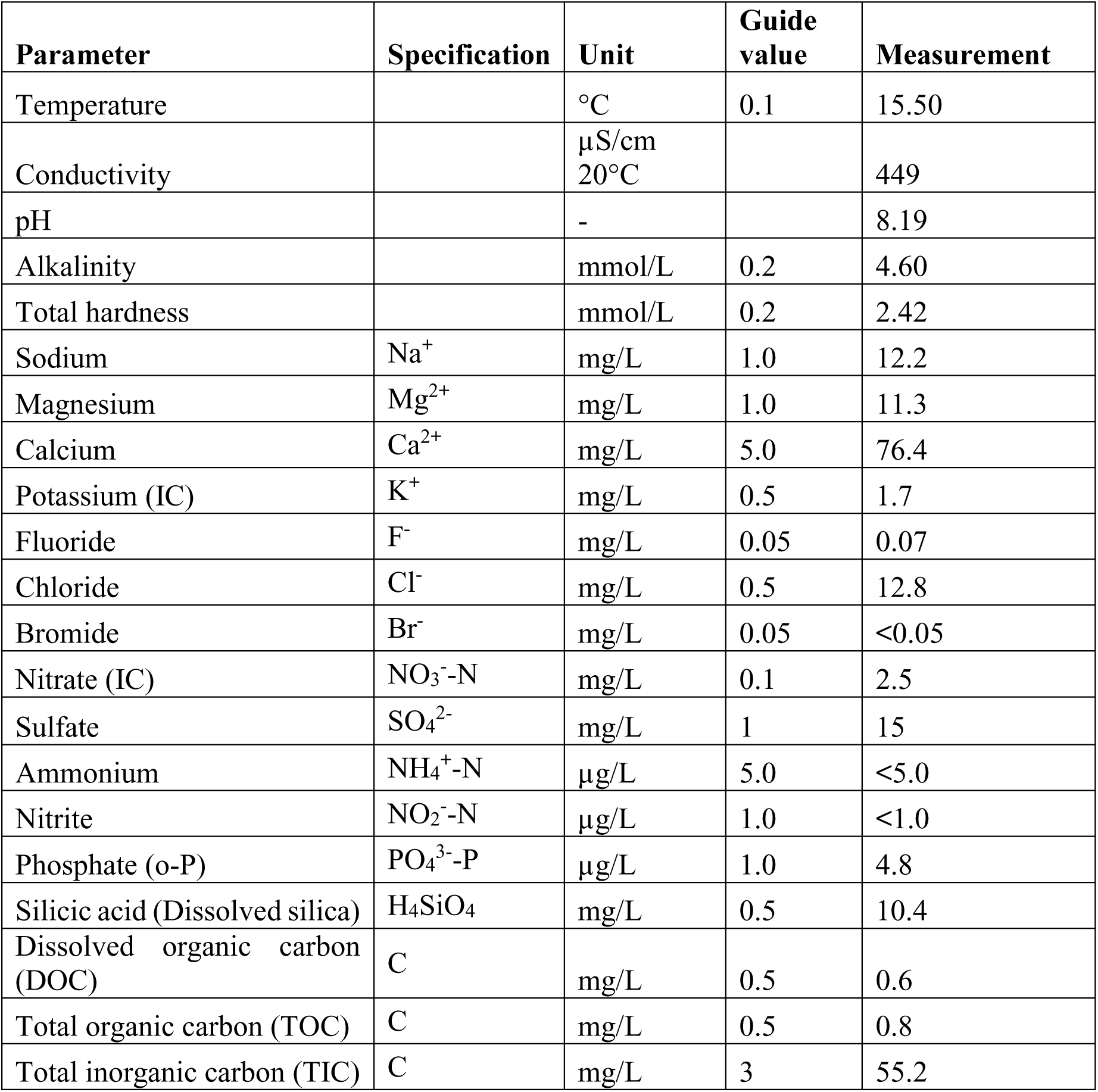
Water matrix characteristics of the water in mesocosm tanks.

**Figure S1:**
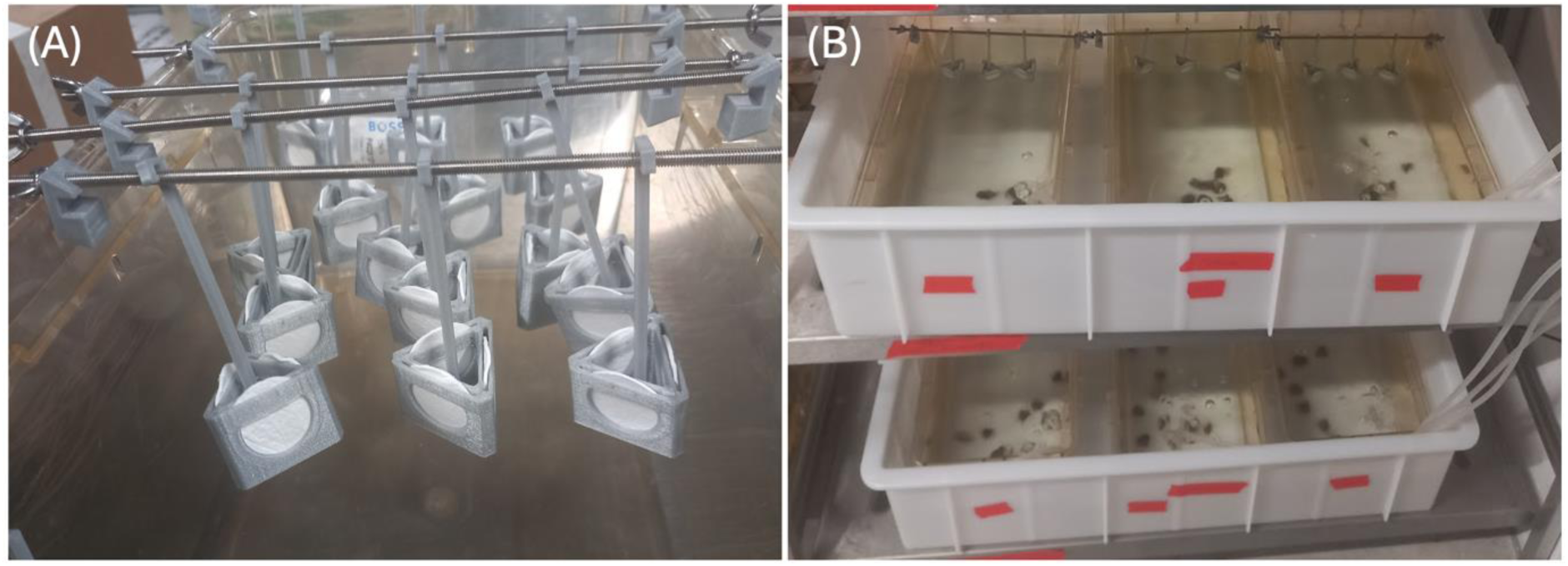
(A) 3D-printed passive sampler housings prepared for deployment in the accumulation dynamics experiments. (B) Passive samplers deployed in triplicate mesocosms during the accumulation dynamics experiments. For 3D-printed housing design, see Schadewell et al., (2026).

**Table S3:**
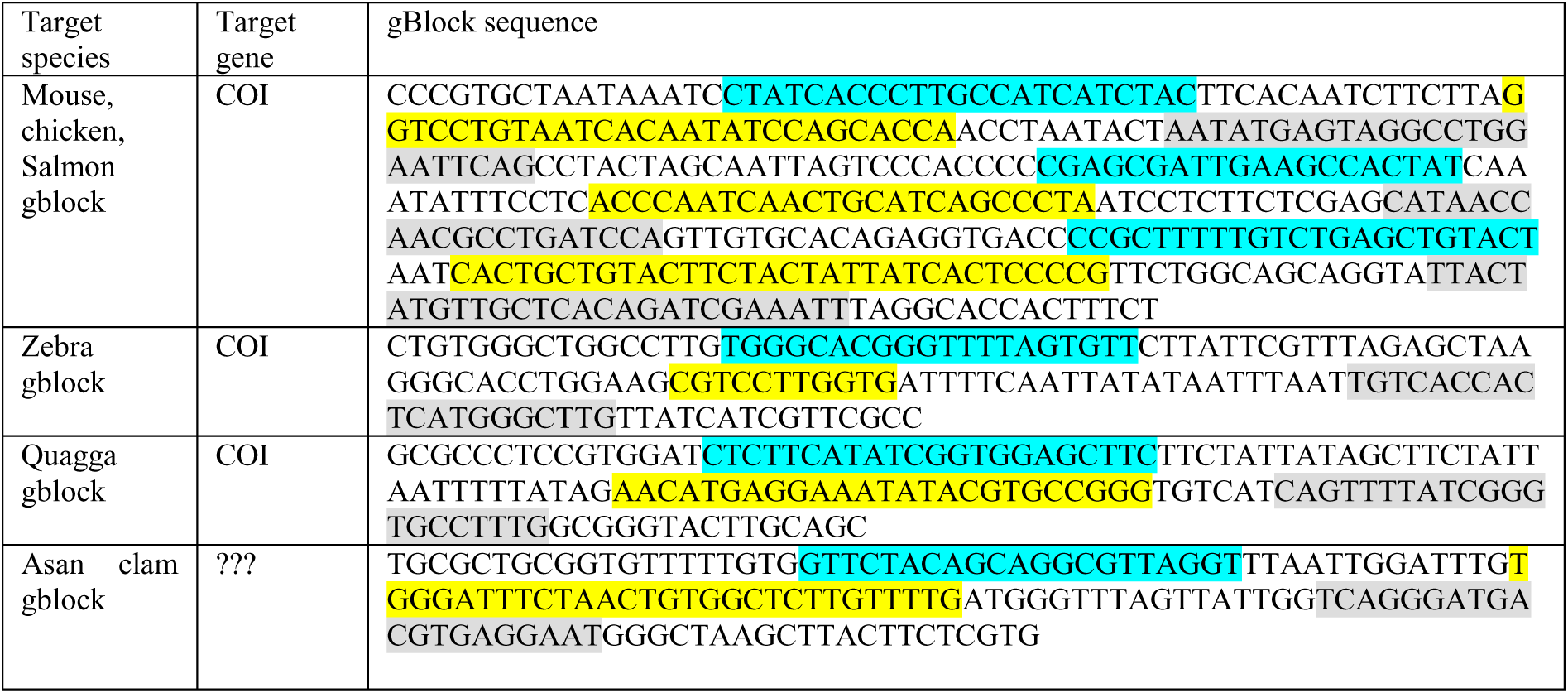
gBlock sequences used in this study. Position of Primers and Probes are marked as follows: Forward Primer, Probe (or probe reverse complement in the case of Quagga), Reverse Primer (reverse compliment).

**Table S4:** DNA concentrations of three species used for mesocosm spike-in experiments. Reported values include the concentration of the spike solution (copies mL⁻¹) and the corresponding expected concentration in mesocosm tank water following dilution.

| Species | DNA concentration (copies/ml spike) | DNA concentration (copies/ml mesocosm tank water) |
| --- | --- | --- |
| Chum salmon | 1.75E+7 | 2,690 |
| Mouse | 2.84E+8 | 43,623 |
| Chicken | 5.89E+9 | 905,620 |

## Notes

### Competing Interest Statement

The authors have declared no competing interest.

### Summary of Updates

The manuscript was revised to include the Acknowledgements section.

